# Vocal correlates of arousal in bottlenose dolphins (*Tursiops* spp.) in human care

**DOI:** 10.1101/2021.04.19.440425

**Authors:** Rachel Probert, Anna Bastian, Simon H. Elwen, Bridget S. James, Tess Gridley

## Abstract

Human-controlled regimes can entrain behavioural responses and may impact animal welfare. Therefore, understanding the impact of schedules on animal behaviour can be a valuable tool to improve welfare, however information on overnight behaviour and behaviour in the absence of husbandry staff remains rare. Bottlenose dolphins (*Tursiops* spp.) are highly social marine mammals and the most common cetacean found in captivity. They communicate using frequency modulated signature whistles which are individually distinctive and used as a contact call. We investigated the vocal behaviour of ten dolphins housed in three social groups at uShaka Sea World dolphinarium to determine how acoustic behavioural patterns link to dolphinarium routines. Investigation focused on overnight behaviour, housing decisions, weekly patterns, and transitional periods between presence and absence of husbandry staff. Recordings were made from 17h00 – 07h00 over 24 nights, spanning May to August 2018. Whistle production rate decreased soon after husbandry staff left the facility, was low over night, and increased upon arrival. Results indicated elevate arousal states associated with the morning arrival and presence of husbandry staff and heightened excitement associated with feeding. Housing in pool configurations which limited visual contact between certain groups were characterised by lower vocal production rates. Production of signature whistles was greater over the weekends compared to weekdays however total whistle production did not differ between weekends and weekdays. Heightened arousal associated with staff arrival was reflected in the structural characteristics of signature whistles, particularly maximum frequency, frequency range and number of whistle loops. Overall, these results revealed a link between scheduled activity and associated behavioural responses, which can be used as a baseline for future welfare monitoring where changes in normal behaviour may reflect shifts in welfare state.

## Introduction

Understanding and monitoring behaviour is a useful tool for welfare assessment [1] as abnormal behaviour may be indicative of poorer welfare [2]. However, the behavioural cues the animal gives must be correctly recognised and interpreted by the observer. Animals housed in captive facilities tend to have structured daily schedules of events such as food provision or routine cleaning that are highly predictable through human-driven cues [3]. Patterns of animal behaviour such as increased alertness [4] and increased vocalisation rates [3] may precede regular events [5] Conversely, reduced activity of the animals has also been documented preceding regular activities such as feeding [6] and public display [4].

Arousal can be classified according to its level (high or low) and its valence (positive or negative) [7,8]. Vocal expression of arousal is a common feature of communication in both humans and non-human animals [9-12] and can reflect responses to immediate experiences [13]. Understanding the vocal cues which act as markers of emotion in mammals [13-15] may be used as a non-invasive tool to monitor welfare for animals held in human care [16-18]. In captive animals, both positive and negative associations with human care givers are observed [19,20] as animals learn to associate humans with rewards and fear through both classical and operant conditioning [21]. Human presence and handling can improve welfare of captive animals, for example in weanling pigs [22] and beef calves [19]. However, in some livestock and poultry, human presence can induce fear and have a negative impact such as decreased growth rates [20,23]. Responses to human presence associated with feeding are often positive [13,24], and the combination of feeding and handling has been shown to play an important role in the development of positive human-animal interactions [25]. Feeding therefore seems to be a suitable candidate to study the behavioural response of a species to a positive interaction.

Animals in captivity are often cared for according to fixed daily schedules where these, as well as other activities with positive outcomes, can elicit behavioural responses for example in the form of anticipatory behaviours [26]. Anticipatory behaviour is a response evoked by reward and is typically linked to elevated arousal states [24], however behaviour has been characterised as either an increase [e.g., 27,28] or decrease [e.g., 29,30] in activity prior to the reward and has been best documented in domesticated mammals [28, 31, 32]. Responses such as anticipatory behaviour can be used as an indicator of animal welfare through measuring the frequency of certain behaviours [33].

Vocalisations often carry prosodic cues about the arousal state of the sender [10] which makes the analysis of vocalisations a suitable tool to assess arousal states. Dolphins live in groups with many social interactions, with vocalisations being their primary method of communication with and between groups. The vocal repertoire of bottlenose dolphins consists of a range of pulsed and tonal sounds [34]. Tonal sounds include narrow-band, frequency modulated whistles to communicate during social interactions [35, 36]. Bottlenose dolphins use ‘signature whistles’ to remain in contact [37-39] and address one another [40]. Signature whistles are individually distinctive whistle types that encode identity information within their frequency modulation pattern and are the most frequently emitted stereotyped whistle type produced by an individual [41]. Although the whistle contour remains stable over time, shifts in whistle production rate, as well as frequency and duration characteristics, may reflect underlying arousal of individual dolphins [38, 42, 43]. As might be expected for a cohesion call, individual production rates of signature whistles increase when motivation to maintain contact or return to the group is strong, i.e., during separation [39] and isolation events [42].

During the day, bottlenose dolphins housed in human care undergo multiple predictable human-controlled events such as feeding, public presentations, training sessions and medical examinations, making them good models for studying patterns in behaviour [44]. Predictable events in captive facilities can evoke behavioural patterns in bottlenose dolphins [4] such as an increase in spy-hopping frequency and surface-looking before any human-animal interaction, including receiving of toys [45]. Vocal behaviour has been reported to be highest when husbandry staff are present and during feeding/training sessions, and lowest at night when numbers of caretakers are reduced [46] with peaks in activity overnight likely associated with bouts of social activity [47]. The morning arrival of caretakers has also been associated with increased vocalisation rate in captive killer whales [48].

As vocal behaviour indicates underlying emotional states, the link between behavioural responses and daily events at a dolphinarium could provide insight into the animals’ life in human care. Additionally, this will provide insight into emotional responses and acoustic cues of emotion in non-human animals. Therefore, the aim of this study was to investigate the overnight behaviour of bottlenose dolphins with a focus on patterns in detections and rates of whistling in response to early morning regimes including husbandry staff presence and feeding. We monitored the vocal response to scheduled events of the population as a whole, as well as at an individual level as animal personalities may vary.

## Methods

This study took place at uShaka Sea World which was established in 2004 and is located in Durban, South Africa. The dolphinarium consists of a covered and open-air pool network with seven interconnected pools of varying size with a combined volume of 11 000 m^3^. At the time of data collection in 2018, the dolphinarium housed ten bottlenose dolphins: three (two male and one female) common bottlenose dolphins (*T. truncatus*), one female Indo-Pacific bottlenose dolphin (*T. aduncus*) and six (four female and two male) hybrids of the two species [49]. The individuals were held in three social groups (see Table 1 for more information on the individuals and social groups) separated by gates which allowed partial visual and full acoustic contact, but not free movement between social groups. All seven pools were utilised by the dolphins during the course of the study, with the number and configuration of pools within which social groups were housed varying throughout the days and overnight. We included pool configuration as a potential predictor variable since visual contact between the different social groups may affect arousal states. Each pool configuration was defined based on which social group was housed in the outdoor presentation pool; configuration 1 – all groups housed in the inside pools with one group having access to the outer interaction pool, and none having access to the outside presentation pool; configuration 2 – the female group housed in outside presentation pool while the mixed and male groups housed in the inside pools with one group having access to the outer interaction pool; configuration 3 – the male group housed in outside presentation pool while the mixed and female groups housed in the inside pools with one group having access to the outer interaction pool; and configuration 4 – the mixed group housed in outside presentation pool while the male and female groups housed in the inside pools with one group having access to the outer interaction pool. The outer interaction pool is directly linked to the outside presentation pool, allowing the dolphins in each of these pools to be in visual contact through the gate. As potential arousal-eliciting activities, we noted morning activities at the dolphinarium which include arrival of husbandry staff (hereafter referred to as ‘staff’) and food preparation at 05h00 as well as feeding and vitamin administration at 06h00. The last public presentation occurs from 15h00 to 15h30 and the last trainer leaves between 17h00 and 18h00, between which times all enrichment devices are removed from the pools.

**Table 1.** Genetic, social grouping and individual data for the dolphins at uShaka Sea World in 2018 (adapted from Gridley et al., 2018)

| Social group | Name | Species | Sex | Capture date | Parents | Signature whistle ID |
| --- | --- | --- | --- | --- | --- | --- |
| Female group | Affrika | *Tt* | F | Captive born | P1 + *Tt* | F1 |
|  | Zulu | *Ta-Tt* Hybrid | F | Captive born | P1 + P2 | F2 |
|  | Khanya | *Ta-Tt* F<sub>2</sub> | F | Captive born | M3 + hybrid | F3 |
|  | Tombi | *Ta-Tt* Hybrid | F | Captive born | P1 + P2 | F4 |
|  | Khethiwe | *Ta-Tt* Hybrid | F | Captive born | P1 + P2 | F5 |
| Male group | Ingelosi | *Ta-Tt* Hybrid | M | Captive born | P1 + P2 | M1/M2 |
|  | Khwezi | *Ta-Tt* Hybrid | M | Captive born | P1 + P2 | M1/M2 |
|  | Kelpie | *Tt* | M | Captive born | Unknown *Tt* x *Tt* | M3 |
| Mixed group | Gambit | *Tt* | M | 08/12/1976 | Unknown *Tt* x *Tt* | P1 |
|  | Frodo | *Ta* | F | 26/06/1979 | Unknown *Ta* x *Ta* | P2 |

### Generating a signature whistle catalogue

The vocal behaviour of individuals was assessed through analysis of signature whistles. To determine the signature whistle for each dolphin, a catalogue was compiled using data collected during temporary isolation sessions in November 2016. Whistle contours are characterised by their time-frequency modulation patterns and in bottlenose dolphins, signature whistles can consist of a single contour or repeated contour (loop). A repeated contour, or multiloop whistle, is either connected where there are no breaks in the entire whistle contour or disconnected with a maximum inter-loop-interval of 0.25 seconds [43].

Acoustic data were collected using one to three High-Tec HTI-96 MIN (flat frequency response of 2 Hz – 30 kHz ± 1 dB) dipping hydrophone(s) attached to a Tascam digital recorder (model DR-680) which sampled the data at 96 000 Hz. Simultaneous vocal notes of observed behavioural data were recorded through a separate headset microphone. Time-frequency spectrograms of acoustic recordings were analysed in Adobe Audition CC v 6.0 (FFT = 1024, frequency range = 0 - 60 kHz, time series window = 10 seconds, Hann window, 50% overlap). To identify the signature whistle of each dolphin, they were temporarily separated with one individual being kept in solitary in one pool between 10 and 20 minutes. The signature whistle was defined as the most common whistle recorded during the temporary isolations and matched to that individual by comparing the relative amplitude of signals on the three hydrophones at different sites. Signature whistles were confirmed using the SIGID bout analysis method [41] where stereotyped calls (at least 3 out of 4) occur in a bout separated by 1 – 10 seconds are considered highly likely to be signature whistles of an individual.

### Investigating patterns of whistling behaviour

Acoustic data were collected over four periods in May, July and August 2018 using a single Sound Trap 300 HF hydrophone (Ocean Instruments, New Zealand, frequency response: 20 Hz – 150 kHz ± 3 dB, sensitivity: 121 dB re. 1 μPa) sampling the data at 576 000 Hz. The hydrophone was placed in the ‘link channel’, an area central to the pool network to which the dolphins did not have access to but was within acoustic range of dolphins held in all pools.

The hydrophone was attached to a 1 kg dive weight and suspended from a rope mid water (at 1.5 m depth, channel depth 2.5 m) held taut by attaching the rope to the roof with a carabiner clip. Ropes tied to the pool sides and roof were used to prevent movement which could produce unnecessary noise on the hydrophone. Acoustic recording commenced in the late afternoon after the final public presentation (between 15h30 and 17h00) and continued until 07h00 the following day. Recording was continuous, but files were restricted to standard 15-minute durations which constrained file sizes.

The first 30 minutes of each overnight deployment was discarded to obtain data unbiased by potential novelty effect as the dolphins may respond to the recorder’s presence in the water. Thereafter, one 15-minute file was selected to represent each hour from 17h00 to 07h00, using the recording which spanned the start of the hour. Each of the selected files were analysed in Adobe Audition CC v 6.0 by visually locating whistle contours in the spectrogram display (FFT = 1024, frequency range = 0 – 60 kHz, time series window = 10 seconds, Hann window, 50% overlap). Each whistle was documented in a database and the signal to noise ratio (SNR) was visually assessed using the following criteria: SNR – 1 whistle is faint and barely visible, SNR 2 – whistle is clear and unambiguous, and SNR 3 – whistle is prominent [50]. Whistles of good quality (those graded as SNR 2 or 3) were either matched to the established signature whistle catalogue and categorised as a signature whistle from a specific individual or categorised as ‘variable whistles’ containing all non-signature whistles from various individuals. The category ‘unclassified whistles’ was used for poor quality whistles (those graded as SNR 1) which did not match to the signature whistle catalogue (Table 2).

**Table 2.** Whistle categories included in the analyses and recording quality rating based on signal to noise assessment.

| Whistle type | Description | SNR |
| --- | --- | --- |
| Signature whistle | A whistle type unique to an individual that matches the signature whistle catalogue | 2 & 3, but can include whistles with SNR 1 if we were confident that they are signatures |
| Signature whistle matching | Matching of an individual's signature whistle by another individual identified through overlapping of whistle contours | 2 & 3 |
| Variable whistle | Loud, unmasked whistles which do not match the signature whistle catalogue | 2 & 3 |
| Unclassified whistle | Masked or partial whistles that were unidentifiable and faint | 1 |

In both the wild and captive facilities, bottlenose dolphins can copy the whistles of others. Such copies can be used in addressing or matching interactions, drawing attention from, or directing information to a particular individual [40, 51, 52]. In some cases, the whistle match might not be an exact replication, but integrate features of the owner’s whistle type or voice [52], however in other cases matches may be indistinguishable from the original. Whistle matching is difficult to identify unless they are overlapping in time. In general, where two whistles with the same contour overlapped in time, we followed [53] in assigning the second whistle as the match and removed it from the signature whistle analyses. However, through the analysis process we identified stereotyped matching behaviour distinct from all other acoustic behaviour observed within the recordings whereby which led to the exclusion of various whistles from the analysis. We noted that the contour of one individual (P2) was emitted at various frequencies in prolonged series of matching interactions. For these stereotyped acoustic interactions, we could not confidently assign whistle production to P2, therefore we termed these whistle sequences ‘square copies’. These square copies, as well as all whistles 30 seconds before and after each event, were removed from the statistical analyses to prevent over-representation of individual P2. All other whistle matching remained in the dataset for the analyses including all whistle types.

To assess the production rate of whistles as a proxy of arousal, we counted the number of whistles in each category (Table 2) per 15-minute recording. We investigated how the production rate of whistles (meaning all whistle types in Table 2 or only signature whistles) was affected by several covariates using a generalised additive mixed model (GAMM) approach using the package ‘gamm4’ version 0.2-6 [54] in RStudio version 4.0.3 [55]. This approach allows fitting of non-linear relationships between variables as well as the inclusion of both fixed and random effects. Whistle production was investigated in terms of both presence/absence and production rate (whistles per minute). A total of four models were built (two for all whistle types in Table 2, and two for signature whistles only) and for each, a variation of the following covariates were tested: ‘hour’ (hour of the day/night), ‘pool configuration’, ‘time of week’ (week vs weekend) and ‘presence of signature whistles’.

Random effects for all four models included ‘sampling day’ to account for the variability of whistles production between sampling days. The selection of covariates was based on testing the assumption that patterns of activities and housing configuration in the dolphinarium lead different levels of excitement which would reflect in dolphin vocal behaviour. Of most interest were the activities of the dolphinarium which occur at scheduled times therefore ‘hour’ was of particular interest. Time of the week was of interest due to the increase of visitors to uShaka Sea World over the weekend. Codes were assigned to working days, ‘week’, (1), which consisted of Monday to Friday, and weekend days, ‘weekend’, (2), which consisted of Saturday and Sunday. The covariate ‘pool configuration’ also contained coded subcategories which were based on which social group was housed in the outdoor presentation pool overnight. All models were run using data between the hours spanning 17h00 to 07h00, with one extreme outlier removed from the total whistle and signature whistle production rate models.

We did not incorporate signature whistle ID as a covariate in the models of the full data series because the data were zero-inflated data due to a lack of whistles emitted in most hours overnight. These models therefore investigate group, but not individual, calling behaviour for whistles. However, individual differences in vocal behaviour are of interest and relevant for animal welfare. Therefore, a separate analysis was conducted to investigate individual differences between 04h00 and 06h00 (04h00 – one hour before staff arrival and food preparation; 05h00 – staff arrival and food preparation; 06h00 – feeding time) when a sufficient number of calls for individual dolphins were available. Data were not normally distributed and had unequal variances (Shapiro-Wilks test and Levene’s test, respectively) even after transformations. Thus, Kruskal-Wallis tests were used to determine if certain individuals significantly increased their signature whistle production rate in response to staff arrival and feeding time. Multiple Dunn *post hoc* tests followed a significant Kruskal-Wallis result. We adjusted the alpha value using the Benjamini-Hochberg method to limit an increase in type I error rate, which is caused by alpha-inflation due to multiple pairwise comparisons.

In addition to changes in whistle production rate, the underlying arousal state may be reflected in the structural parameters of whistles such as duration and frequency characteristics [11, 42]. We investigated this for each individual by selecting five standard whistle parameters from signature whistles namely minimum and maximum frequency, frequency range, duration and number of loops, as well as production rate for each signature whistle. Measurements were taken from time-frequency spectrograms (FFT = 1024, frequency range = 0 – 40 kHz, time series window = 5 seconds, Hann window) of the fundamental frequency of 33 – 392 signature whistles from each animal in Raven Pro v1.6.1 [56]. For each signature whistle, production rate data were pooled into time periods representing staff absent (20h00 – 04h00) and staff present (05h00 – 07h00). Thereafter, production rates per hour were recalculated for each time period. Times 17h00 – 19h00 were omitted from this analysis due to the uncertainty of daily staff presence that occurred at these times. Data were not normally distributed and had unequal variances (Shapiro-Wilks test and Levene’s test, respectively) therefore paired-sample Wilcoxon tests were used to investigate change in production rate between the pooled time. Additionally, all five signature whistle parameters were individually compared between the two pooled time periods for six of the ten individuals with sufficient data (at least five signature whistles for each individual for each pooled time period). Again, data were not normally distributed and had unequal variances (Shapiro-Wilks test and Levene’s test, respectively), therefore Wilcoxon tests were used to investigate change in the acoustic parameters of signature whistles between times of ‘staff absent’ and ‘staff present’.

## Results

A total of ten signature whistles were documented for the group of ten dolphins, of which eight were assigned to individuals (see Table 1, Fig 1A). The two remaining signature whistles could not be confidently differentiated between individuals M1 and M2, which are hybrid siblings which engage in a significant amount of whistly matching. All square whistle copying sequences were removed from the analyses (Fig 1B).

**Fig 1.**
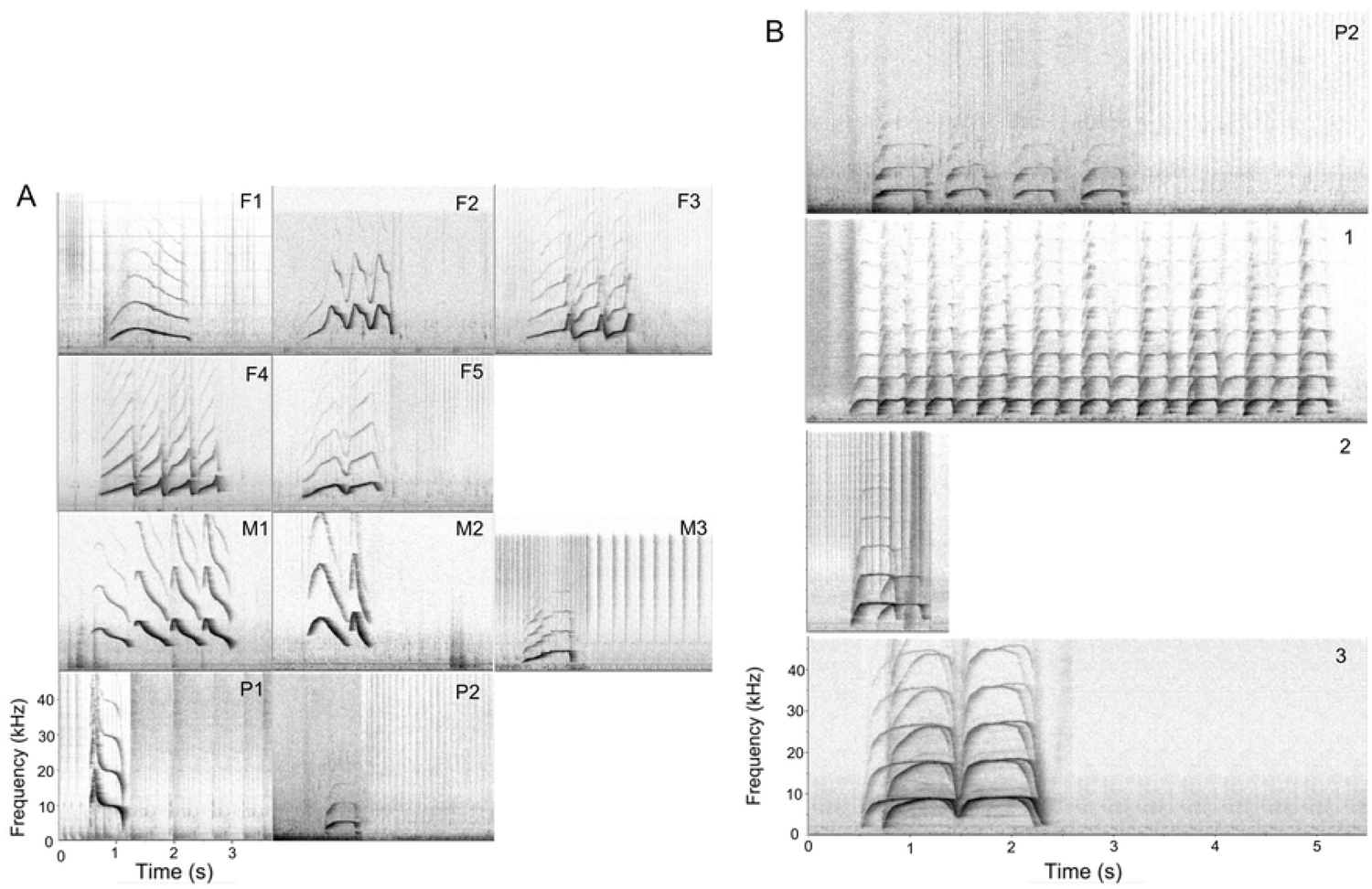
Whistle catalogues. (A) Signature whistle catalogue of all ten dolphins housed at uShaka Sea World in 2018 (F1, M3, P1 and P2 are single loop whistles; F2, F3 and F5 are connected multiloop whistles; F4, M1 and M2 are disconnected multiloop whistles). (B) Examples of (1) low, (2) middle and (3) high frequency square whistle copying, with a similar contour shape to dolphin P2.

Nocturnal whistling behaviour was investigated from acoustic recordings made over 24 sampling nights (17 on weekdays and 7 on weekend days) which were conducted in four hydrophone deployment periods ranging from five to seven nights in duration. From this, 88 hours of acoustic data were analysed with a total of 2640 documented whistles, 1647 of which were signature whistles (62.4%), 168 variable whistles (6.3%), 569 unclassified whistles (21.6%), 3 signature whistle matching (0.1%), 98 square whistles (3.7%) and 155 square whistle copying (5.9%) (Table 2). Square whistle copying behaviour was uncommon and occurred sporadically with no particular temporal trend (S1 Fig). Furthermore, these square copying events occurred predominantly during the night when overall signature whistle production was at a minimum. In addition to P2 not confidently being assigned a whistle in these copying interactions, a closer inspection of these square whistle copying events indicated that the vocal behaviour was more involved than a simple owner-copy interaction. In the 30 seconds before and after these whistle copying events that occurred throughout the data set, whistles were produced most by three of the animals (S2 Fig). After removing all whistles 30 seconds before and after each square whistle event, 2281 whistles, 1561 of which were signature whistles, were included in the analyses.

The GAMMs show a temporal trend in presence and production rate of whistles with two clear peaks, one in the late afternoon and one in the morning (Fig 2; Table 3 – Models 1 to 4; variable ‘hour’; *p* < 0.001). These peaks correspond to staff presence, particularly to the time of arrival of staff and preparation of food in the morning (05h00 – 06h00). Pool configuration also influenced overall whistle presence and production rate. Total whistle presence and signature whistle presence significantly decreased when dolphins were housed in pool configuration 2 (female group housed outside) compared to configuration 1 where all animals were housed in the indoor pools (Fig 3A; Table 3 – Model 1 and 3, variable ‘pool configuration’; *p* < 0.05). Total whistle production rate decreased when dolphins were housed in pool configuration 4 (mixed group housed outside) and signature whistle production rate decreased in pool configuration 3 (male group housed outside) in comparison to configuration 1 (Fig 3B; Table 3 – Model 2 and 4, variable ‘pool configuration’; *p* < 0.05).

**Fig 2.**
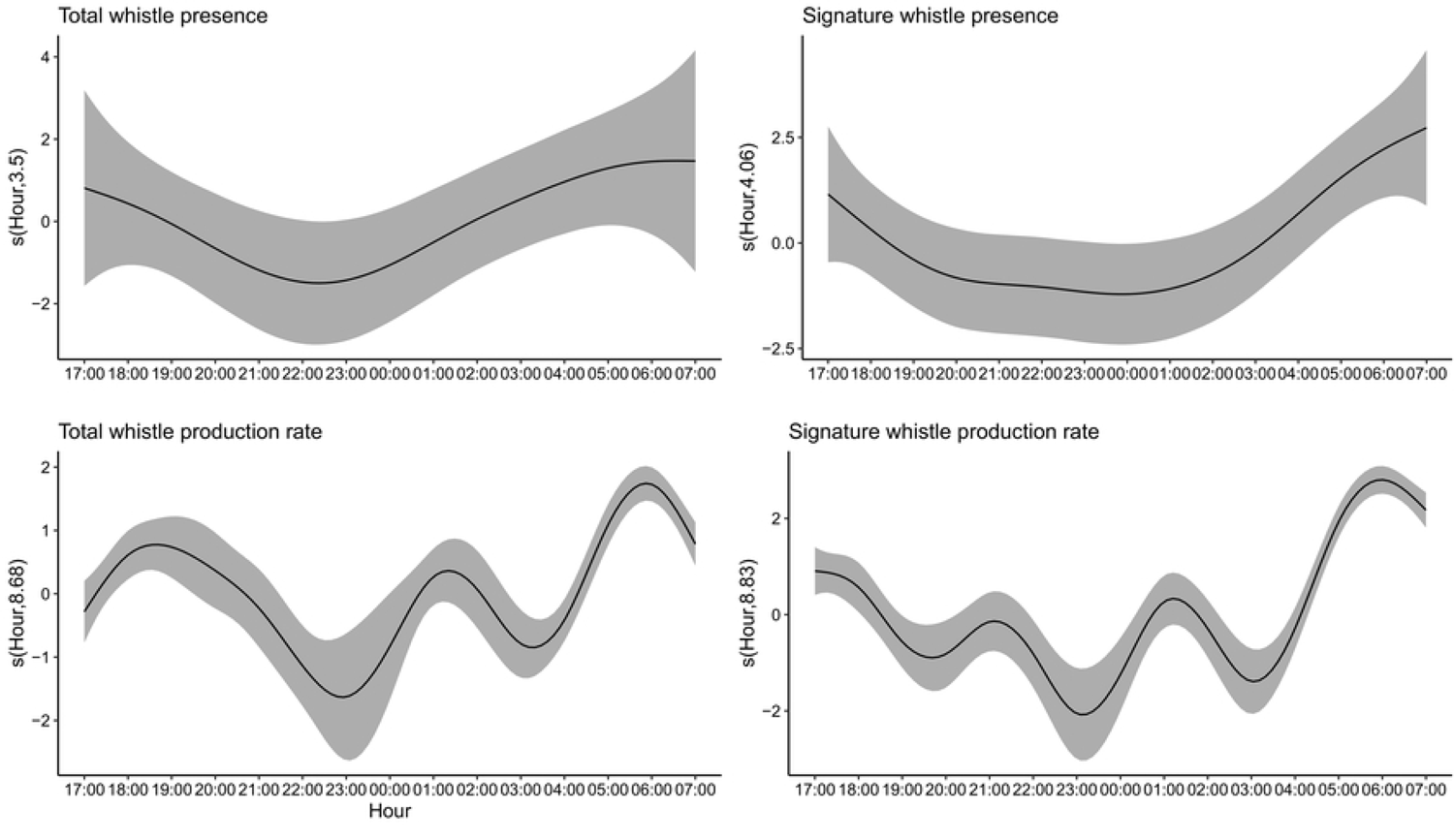
**GAMM summary of smoothing term ‘hour’ for all four models.**

**Fig 3.**
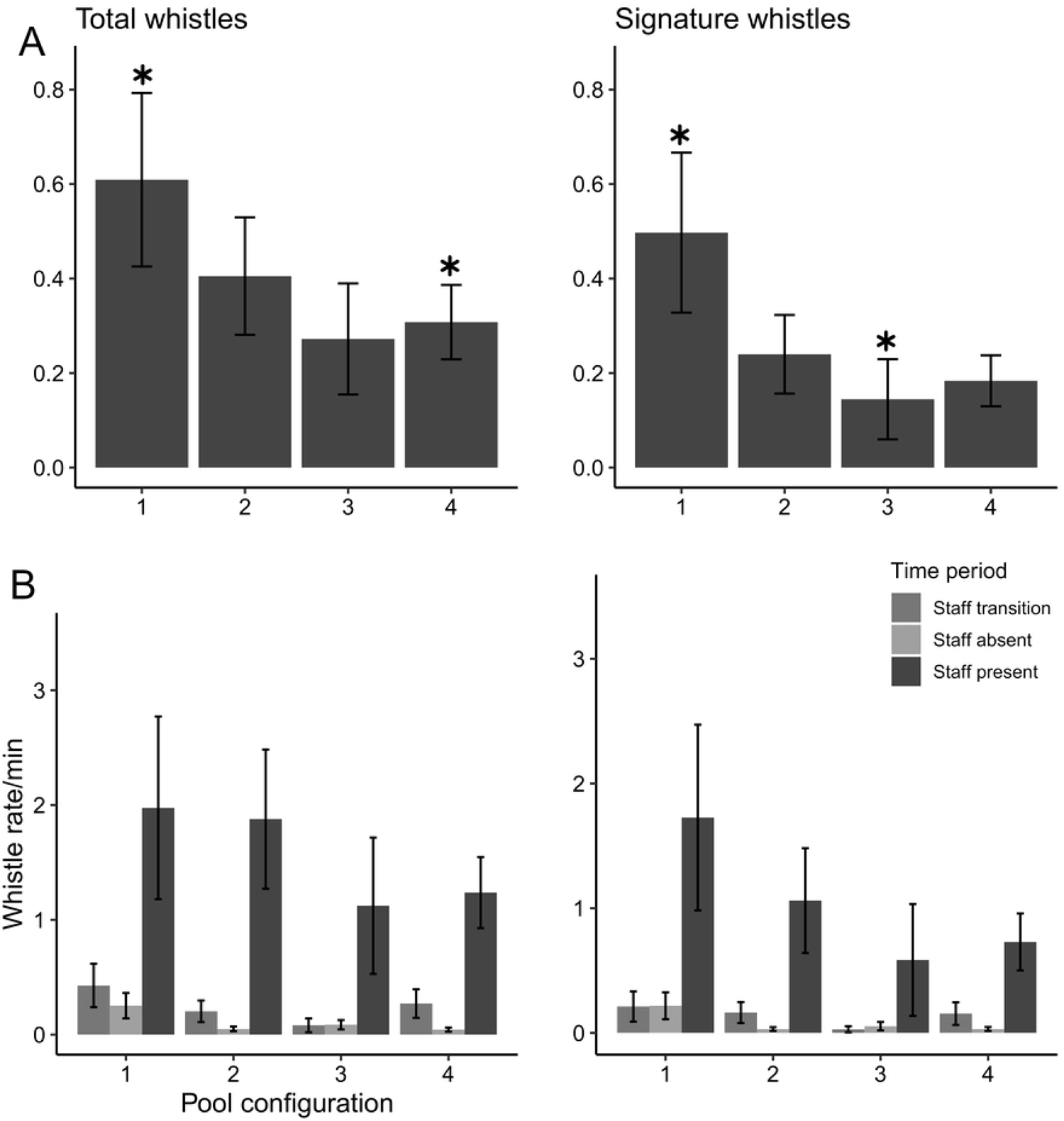
Total whistle and signature whistle production rate compared between pool configurations. (A) Production rates (B) Production rates split into time periods associated with husbandry staff presence.

**Table 3.** GAMM results for all presence/absence and production rate models (only *p-*values of significant results presented)

| Response variable | N | Explanatory variables |  |  |  | Adjusted R-squared |
| --- | --- | --- | --- | --- | --- | --- |
|  |  | Hour | Pool configuration | Time of week | Signature whistle presence |  |
| **Model 1:** | 347 | < 0.001 | 0.042 (Intercept) | NA | - | 0.649 |
| **Whistle presence** |  |  | 0.020 (config 2) |  |  |  |
| **Model 2:** | 346 | < 0.001 | 0.044 (Intercept) | NA | < 0.001 | 0.521 |
| **Whistle count** |  |  | 0.025 (config 4) |  |  |  |
| **Model 3:** | 347 | < 0.001 | 0.034 (Intercept) | NA | NA | 0.279 |
| **Signature whistle presence** |  |  | 0.017 (config 2) |  |  |  |
| **Model 4:** | 346 | < 0.001 | 0.001 (Intercept) | - | NA | 0.233 |
| **Signature whistle count** |  |  | 0.008 (config 3) |  |  |  |

The proportion of whistles produced during ‘staff transition’ (the afternoon period where staff were leaving and/or had just left), ‘staff absent’ and ‘staff present’ indicates that this trend in whistle production driven by pool configuration is prevalent during the overnight period where staff were absent (Fig 3A, B). The likelihood of whistle or signature whistle occurrence per hourly bin did not differ between weekdays and the weekend, however total signature whistle production rate was greater over the weekend (Fig 4; Table 3; Model 4, variable ‘Time of Week’). Although not significant, the inclusion of this variable increased model performance (R-squared from 0.178 to 0.233) which indicates that the time of the week plays a role in the production of signature whistles at the facility. Total whistle presence and production are highly dependent on the presence of signature whistles (Table 3; Model 1 and 2, variable ‘Signature whistle presence’). Although not significant in Model 1, this variable significantly increased overall model performance (R-squared from 0.302 to 0.649). The low-average adjusted R-squared values in all four models (Table 3; adjusted R-squared range = 0.233 – 0.649) indicate that there are other factors influencing whistle and signature whistle production rates that cannot be explained by the covariates in these models.

**Fig 4.**
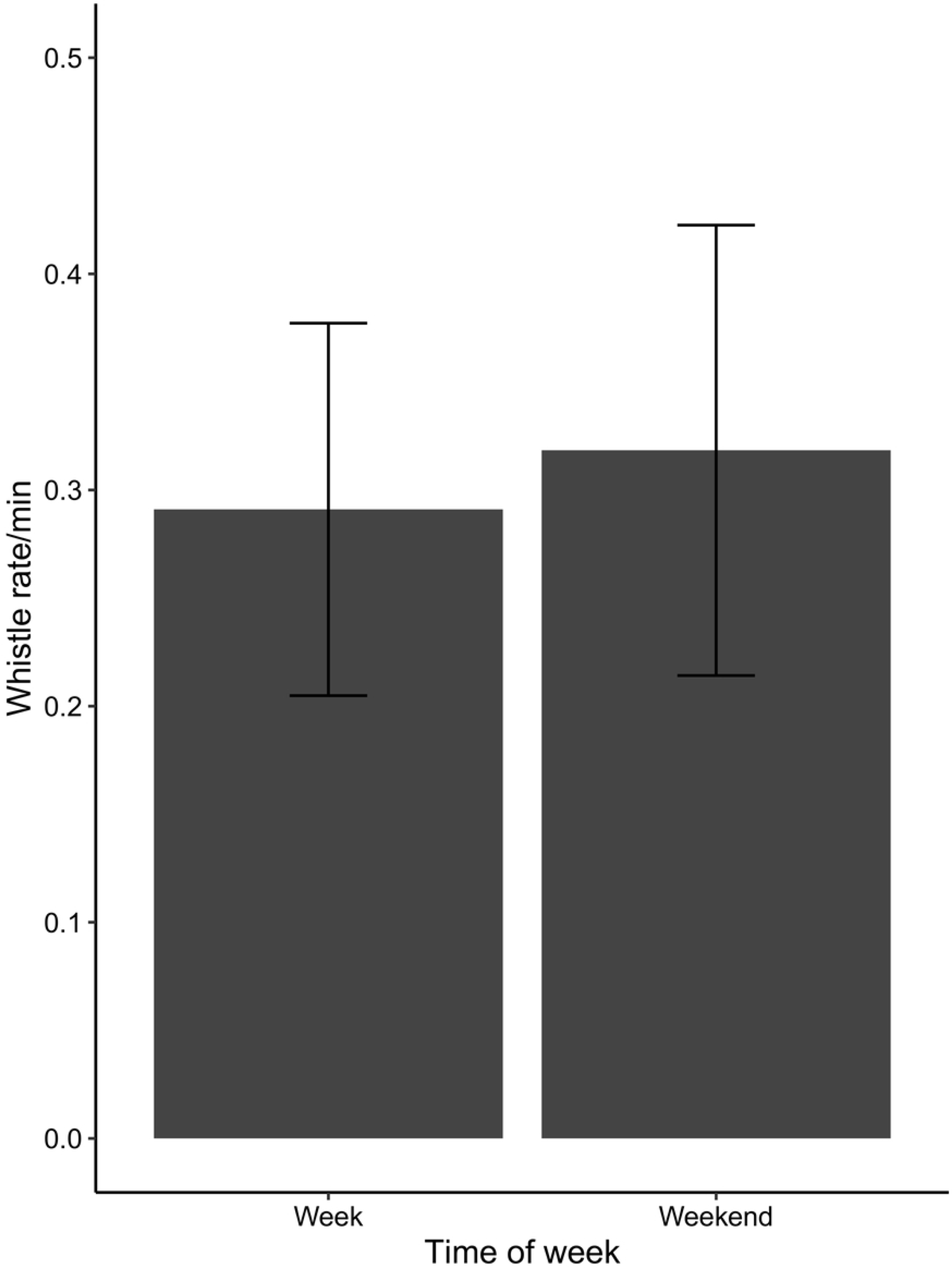
**Signature whistle production rate compared between weekdays and weekend days.**

There was a large difference in signature whistle production between individuals. Of the 1561 signature whistles identified ∼ 60% were produced by three individuals (two female and one male, Fig 5). The three least vocal animals contributing ∼ 7% to the total signature whistle production were all males (Fig 5). A comparison of individual signature whistle production rate was investigated in the morning when total whistle production was at its highest and there was sufficient data (04h00 to 06h00) to determine individual differences in response to staff arrival (05h00) and feeding time (06h00). Kruskal-Wallis tests indicated that the signature whistle production rate of five of the dolphins increased significantly only in response to feeding time (Fig 6A; no significant difference between 04h00 and 05h00, p < 0.05, n = 866). The remaining five individuals did not exhibit a significant increase and P1 produced no whistles during feeding time. Additionally, some of the animals were completely silent overnight when no staff were present. Investigating individual differences in signature whistle production between periods with staff absent (20h00 – 04h00) and with staff present (05h00 – 07h00), F2 was highly vocal during both of these time periods (Fig 6B) and signature whistle production from all dolphins increased with staff present (Fig 6B, p < 0.05 for five of them, n = 1413). In the absence of staff, bouts of whistling occurred sporadically between sample days, driven mostly by F2 as well as four other individuals. Although signature whistle identification could not be included in the above-mentioned models as a covariate, the results strongly suggest that individual differences in signature whistle production are important.

**Fig 5.**
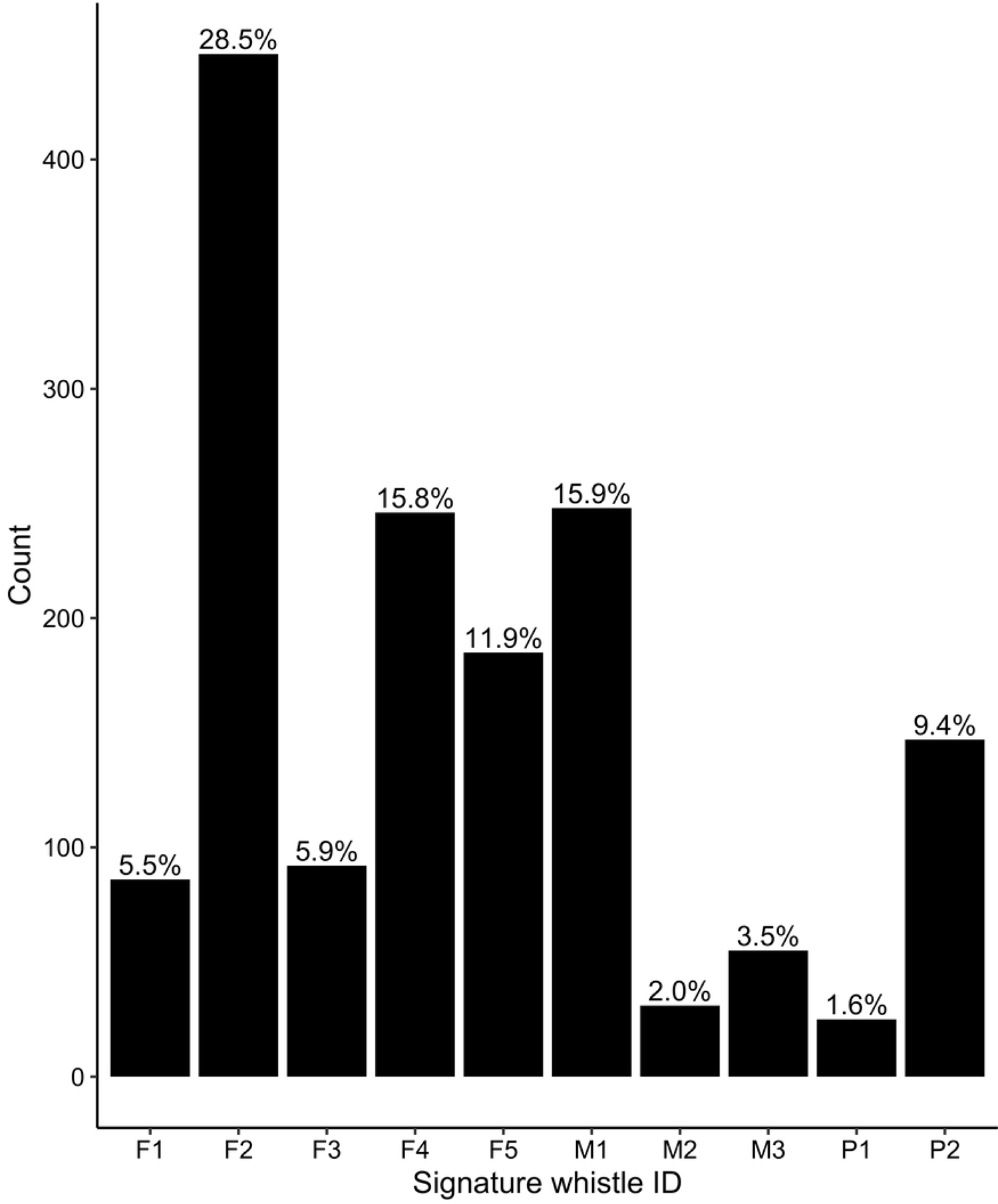
**Total count of signature whistles produced throughout the study period.**

**Fig 6.**
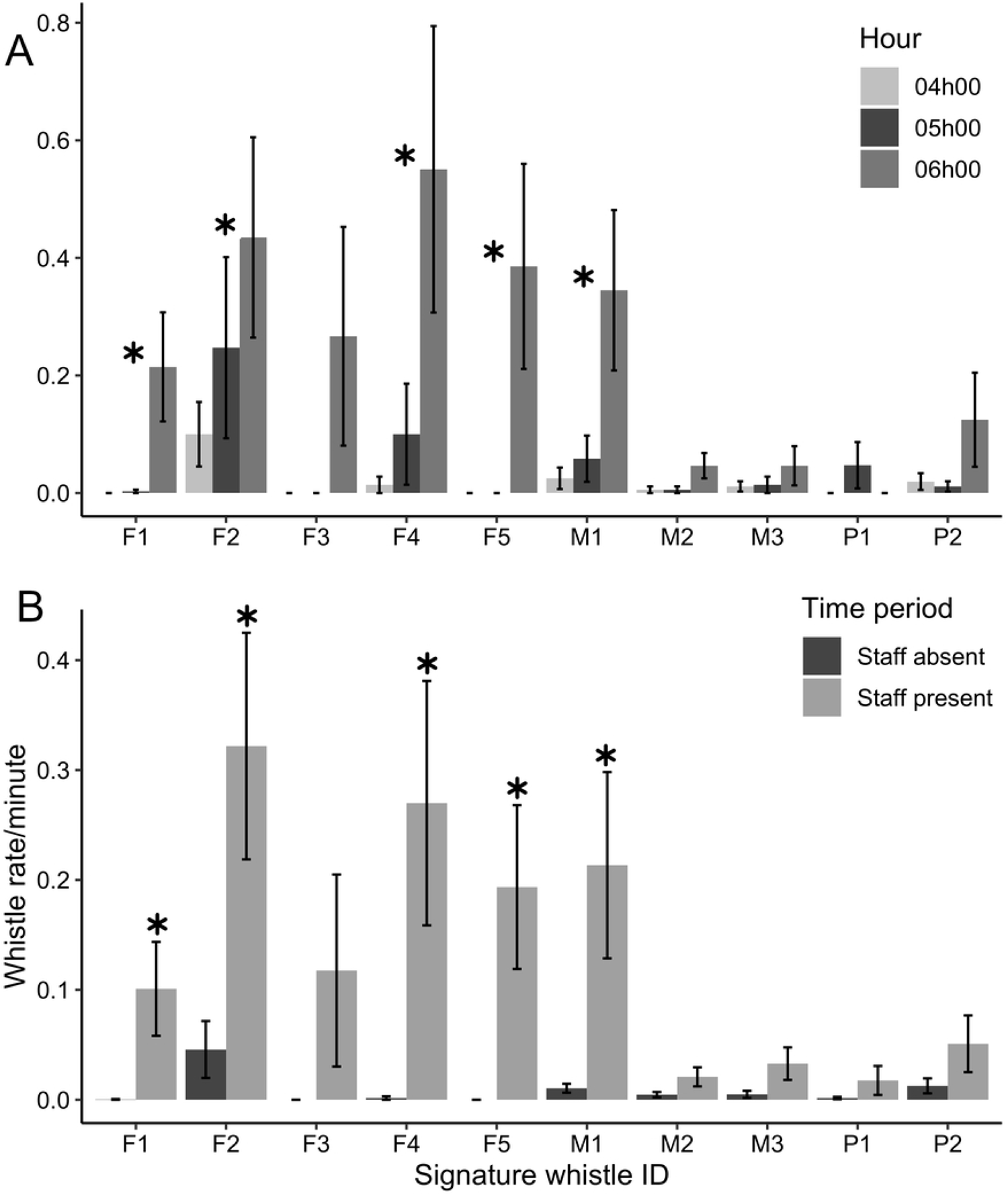
Individual vocal responses to feeding and staff presence. (A) Signature whistle production rate of each individual compared across three hours (04h00 – one hour before staff arrive, 05h00 – staff arrival and food preparation, 06h00 – feeding). Significance indicated by asterisks is consistent among individuals, with no significant shift between 04h00 and 05h00. (B) Whistle production rate per hour of all individuals when staff were absent and when staff were present. Significance between time periods indicated by asterisks.

Matched pairs Wilcoxon tests showed significant differences in the acoustic parameters of their signature whistles between periods of staff absence and presence (Fig 7). The three animals which exhibited the most changes in whistle characteristics all showed a significant decrease in maximum frequency and frequency range when staff were present. The two other individuals did not exhibit any significant shifts in whistle characteristics. Overall, most of the individuals showed significant decreases in maximum frequency (average: from 12.8 to

**Fig 7.**
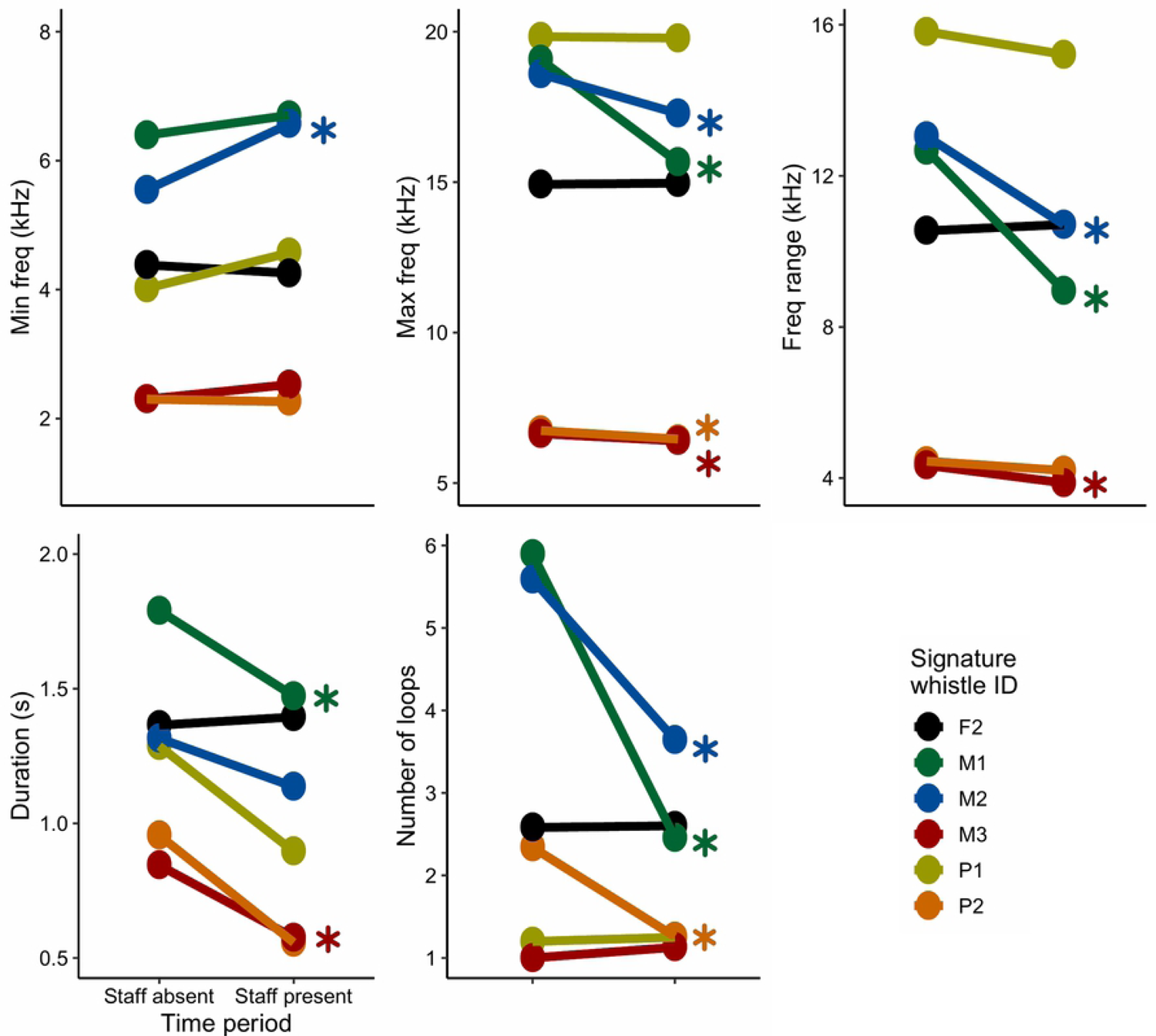
**Shifts in signature whistle parameters in response to staff presence. Asterisks indicate significance.**

11.4 kHz), half of them significant decreases in frequency range (average: from 10 to 7.9 kHz) and number of loops (average: from 4.6 to 2.5 loops per whistle), while two individuals exhibited significant decreases in whistle duration (average: from 1.35 to 1.05 s) and only one individual a significant increase in minimum frequency (from 5.5 to 6.6 kHz).

## Discussion

The overall night-time vocal behaviour between the afternoon and morning peak of whistle production was fairly quiet, with some bouts of whistling behaviour during the night occurring randomly between recording nights. This is in accordance with [57] who found that peaks in nocturnal vocal activity of captive bottlenose dolphins may occur. Sleeping/resting occurs predominantly during the night and accounts for up to 87% of nocturnal behaviour of captive bottlenose dolphins [58]. This explains the decrease in vocal activity during this night-time period, however from this study there has been no indication what might be driving the bouts of social behaviour. It has also been shown that captive bottlenose dolphins increase vocal and social interactions before their rest periods, possibly to promote synchronous swimming, a behaviour observed in wild and captive dolphins [58]. Similarly, in a study of captive bottlenose dolphins (*T. truncatus*) at facilities in Japan, vocal production rates increased during the day when their human caretakers were present [46, 58] and vocal behaviour, including whistle production rate, decreased at night when caretakers were absent [46]. These studies indicate a positive relationship between caretakers and animals in captivity may be beneficial to the welfare of the animals [59].

Daily arousal patterns which coincided with presence of staff and caretakers at the facility, specifically at the end of the day (last daily presentation) and early in the morning (food preparation and feeding), suggest excitement associated with these activities. Excitement is a positive arousal state [8, 60], and is less commonly studied than negative arousal. Vocal responses indicating positive and negative emotional states are hard to differentiate as they may share vocal features [61]. In the current study, signature whistle production rates increased in the morning during staff arrival and food preparation, significantly intensified during feeding and then decreased one hour after feeding and was driven by half of the individuals in the facility. At feeding time, five dolphins significantly increased signature whistle production. In wild bottlenose dolphins an increase in vocal activity is associated with foraging which may be to maintain contact or to recruit individuals [62]. Food-anticipatory activity describes increased arousal before feeding events on strict daily schedules [63] and has been observed in captive animals during scheduled feeding times [46, 64]. The emotional value of anticipatory behaviour is thought to reflect the balance of the reward system experienced by animals before and during feeding [27]. Increased vocalisations around feeding time have also been documented in captive false killer whales (*Pseudorca crassidens*), common bottlenose dolphins (*T. truncatus*) and common dolphins (*Delphinus delphis*), where vocalisation rates increased upon the arrival of their caretakers and was maintained or intensified throughout feeding and decreased immediately after [48, 64]. In wild animals, detection of food may lead to an increased arousal state followed by temporary “elation” after capture of prey [8]. Other studies however observed a decrease in activity and arousal before human-animal interactions including feeding, for dolphins [4] and chimpanzees [6].

Of the four males in this analysis, the three younger males exhibited significant changes in most of the whistle parameters, all including a decrease in frequency range from a period with staff absent to a period with staff present. One male decreased whistle production rate when staff were present and two males exhibited significant decreases in whistle duration, one of which also decreased number of loops. The oldest male did not significantly shift any whistle parameters between these two time periods. Of the two females included in this analysis, F2 did not significantly shift any whistle parameters between these two time periods while the oldest female in the facility decreased maximum frequency and number of loops.

Some dolphins may emit whistle loops faster than normal when they are excited [65] therefore duration is a function of number of loops [42]. Whistle duration has been shown to increase during isolation [42], and much like number of loops, in the context of excitement and stress, whistle duration seems to be the opposite. Whistle frequency parameters in bottlenose dolphins such as production rate and number of whistle loops are closely related to the level of arousal of an individual [42, 43]. The number of loops and whistle production rates vary by context and an increase in these parameters may indicate stress [43] or excitement [66] in bottlenose dolphins. In the current study, the whistle production rate significantly increased in six of the ten individuals, however a significant decrease in the number of loops was only exhibited by three of the six dolphins evaluated. An increase in the number of loops in the context of stress [43] and a decrease in the number of loops in the context of excitement (as in the current study) suggest that loop number may be a useful tool in monitoring positive and negative arousal states in bottlenose dolphins. Although certain shifts in signature whistle parameters appear to be indicative of individual arousal, differences are not consistent across individuals [42]. The same applies to the dolphins at this facility where individuals had different combinations of whistle parameter shifts, therefore the dolphins should be individually considered and monitored.

In this study the dolphins were less vocal when the females were housed in the outdoor presentation pool and produced whistles less frequently when the mixed group was housed outside and produced signature whistles less frequently when the male group was housed outside and were most vocal in pool configuration 1, when they were all housed in the indoor pools (Fig 3A). This is contradictory to what you might expect from a cohesion call [39, 42] and indicates an emotive or social reaction in response to these different types of group separation. Although pool configuration affects overall whistle production rates, a vocal response to staff presence or absence is much less clear (Fig 3B) and somewhat contrary.

When housed inside, human activity is in view of all dolphins, including the arrival of staff, therefore the strongest behavioural response would be expected here. Whistle production rate in response to staff presence is higher in configuration 2 for total whistles, and similar in configurations 1 and 2 for signature whistles, which is not what was expected. The social dynamics are quite different among the three groups housed at uShaka Sea World and although they are housed in separate social groups, they are always in acoustic (and mostly visual) contact with one another. This may allow inter-group social bonds to be formed, which has also been documented in wild meerkats [67] and wild bottlenose dolphins [68].

Bottlenose dolphins live in complex societies and depend greatly on close social bonds [69]. Because of this, they are more likely to suffer from social-related stress in both the wild and captivity [70] however there is great opportunity for these animals in captivity to achieve positive welfare states due to strong social bonds within their social groups [71].

Whistle production rate was higher over the weekend compared to weekdays. While the number of visitors were larger on weekends than during the week, the daily schedule was the same regardless of the day of the week. Visitors to captive facilities play a role in the behavioural responses of mammals, however not all captive mammals are affected by visitor presence and noise [72]. For example, negative responses to visitors in captivity has been documented in nonhuman primates [73-76], whereas captive meerkats are behaviourally unresponsive under different intensities of visitor behaviour [77]. Bottlenose dolphins undergo activities in the form of public presentations which has been documented to elicit positive arousal rather than stress or negative arousal [4]. However, the relationship between the magnitude of arousal proceeding a presentation and audience size has not yet been investigated. Increased whistle production rate over the weekend could not be directly linked to an increase in visitors as data were collected once visitors had left, however residual excitement from large crowds could have possibly been carried over into the overnight recording periods.

Regarding the overall signature whistle production, all square copies were removed from the analysis and in doing this, there was a possibility that signature whistles of P2 were underrepresented. Square whistle copying interactions are important in terms of social interactions and bonds between certain animals and understanding this will give insight into their social vocal behaviour as a whole. Of the ten dolphins in the study, the most vocal during both overnight and morning periods was a female, the third youngest dolphin. The least vocal was a male, the oldest dolphin at the facility. Age has been documented to affect signature whistle production rate in bottlenose dolphins where higher signature whistle rates are present in younger dolphins and decreases with age, more quickly in males [42, 43]. Differences in whistle production in bottlenose dolphins between sexes has previously been documented due to differences in social histories of males and females [78]. Because the whistles of hybrids M1 and M2 could not be differentiated, we could not investigate the role that age and sex play together in vocal activity of these dolphins in more detail.

The most vocal individuals in this study were predominantly hybrids, all of which are offspring of P1 and P2. Differences in behaviour between purebred and hybrid cetaceans has been reported [79]. Hybridisation of captive bottlenose dolphins and other dolphin species has been widely documented [58, 80-82], however hybridisation between bottlenose dolphin species in captivity has seldom been seen [see 49, 83]. Although hybridisation is naturally occurring in 10% of animal species [84], in the wild there is little overlap of home ranges of *Tusriops* spp., limiting inter-species mating opportunities of bottlenose dolphins [49].

Hybridisation has been documented to influence behaviour in animals; for example, neonatal hiding behaviour [85] and vocalisations [86] of hybrid deer species are strongly influenced by genetic history and display intermediate characteristics of the parent species. Similarly, intermediate behaviour of free-ranging porpoise hybrids has been documented [79]. Although not included in the analyses due to small sample sizes, species may play a role in signature whistle production of bottlenose dolphins. Welfare assessments in the past were heavily focussed on monitoring behaviour indicative of poorer welfare as good welfare was thought to result from lack of suffering [87].

Vocalisations in farm animals have been used as a measure of welfare on farms [88]. Animals that express more arousal and anticipation for events or rewards may be in poorer welfare however some level of arousal before a positive event, such as feeding, reflects positive emotions [89]. One such emotion is excitement [8], which is the emotional state that the dolphins in this facility appear to be reflecting when they increase the production rate of social sounds. Additionally, some signature whistle characteristics reflect this excitement associated with the presence of staff at the facility. Undesirable behaviour may only be present in a certain group or sex, which emphasises the limitations of a unified approach to monitoring welfare and the importance to explore individual welfare states [90]. This all provides a useful index; where production rate and whistle characteristics would usually shift within this population, an abnormal change at either a group or individual level could be indicative of a shift in welfare state.

## Acknowledgements

We would like to thank Larry Oellermann and Gabrielle Harris at uShaka Sea World for allowing us to spend time at the dolphinarium to collect data, and trainer Kelly de Klerk and team for assisting with acoustic data collection.

## Supporting information

S1 Fig. Total number of square whistle copying events during each time bin throughout the study period.

S2 Fig. Total whistle counts, as well as P2 as the initiator of the copying events (Kriesell et al., 2014).

